# Effects of footshock stress on social behavior and neuronal activation in the medial prefrontal cortex and amygdala of male and female mice

**DOI:** 10.1101/2021.12.16.473017

**Authors:** Mariia Dorofeikova, Chandrashekhar D Borkar, Katherine Weissmuller, Lydia Smith-Osborne, Samhita Basavanhalli, Erin Bean, Avery Smith, Anh Duong, Alexis Resendez, Jonathan P Fadok

## Abstract

Social behavior is complex and fundamental, and its deficits are common pathological features for several psychiatric disorders including anxiety, depression, and posttraumatic stress disorder. Acute stress may have a negative impact on social behavior, and these effects can vary based on sex. The aim of this study was to explore the effect of two days of footshock stress on the sociability of male and female C57Bl/6J mice measured using a standard social interaction test. Animals were divided into two main groups of footshock exposure or context exposure control. Each group had mice that were treated with either the benzodiazepine alprazolam, or vehicle. In all groups, neuronal activation during social interaction was assessed using immunohistochemistry against the immediate early gene product cFos. Although footshock stress did not significantly alter sociability or latency to approach a social interaction counterpart, it did increase defensive tail-rattling behavior specifically in males. This stress-induced increase in tail-rattling was alleviated by alprazolam, yet alprazolam had no effect on female tail-rattling behavior in the stress group. Alprazolam lowered cFos expression in the medial prefrontal cortex, and social interaction induced sex-dependent differences in cFos activation in the ventromedial intercalated cell clusters. Overall, our results suggest that acute footshock stress induces sex-dependent alterations in defensiveness and patterns of cFos activation during social interaction tests.

## Introduction

Social behavior is important for survival, and social deficits are common pathological features for a variety of mental illnesses including social anxiety disorder, depression, and posttraumatic stress disorder [1, 2]. More women than men suffer from these disorders, yet there is a paucity of data on sex differences in social behavior after stress [3]. There are data suggesting that childhood trauma leads to more aggressive behavior in men and more social withdrawal and avoidance in women [4], and women with social anxiety disorder report greater clinical severity, which may be associated with stressful life experiences [5]. Therefore, understanding sex differences in the effects of traumatic stress on social behavior, as well as the underlying neural substrates that potentially control this behavior, has important translational relevance.

Animal models of stress and trauma show alterations in many aspects of behavior [6]. In general, both acute and chronic stress have been found to lead to social fear and withdrawal in rodents [7]. One common model of traumatic stress in rodents is footshock exposure [8]. However, there is a lack of data investigating whether footshock stress influences mouse social behavior in a sex-dependent manner.

It is also unclear whether there are sex differences in the activation of brain regions responsive to social encounters following acute stress. Several lines of evidence suggest that neural activity in the medial prefrontal cortex (mPFC) is important for social behavior and subregions of the mPFC are differentially activated following social interaction. Neurons in the infralimbic cortex (IL) are preferentially activated in response to social cues compared to neurons in the prelimbic cortex (PL) [9], and a social neural ensemble within the IL may contribute to the social buffering of fear after fear conditioning [10].

Several subnuclei of the amygdala have also been implicated in social behavior. The role of the basolateral amygdala in social interaction following different stress paradigms has been well-established [12–14]. For example, activation of the basolateral amygdala leads to reduced social interaction in a social interaction test [11]. The medial nucleus of the amygdala (MeA) is also involved in both social behaviors and responses to stressors [15]. cFos expression in the intercalated nucleus of the amygdala (ITC) is increased during social buffering in rats [16], and social interaction is impaired in mice with altered migration and differentiation of ITCs [17, 18]. Among the amygdala regions implicated in social behavior, the central nucleus of the amygdala (CeA) is relatively unexplored. There is recent evidence, however, that CeA circuits may be linked to sociability, and some manipulations of CeA activity impact social behavior [18–21].

Thus, multiple brain regions are involved in social cognition [15, 16], but the differences in social approach or avoidance behavior following acute stress are still poorly understood. In the current study, our goal was to assess the extent to which there are sex differences in sociability using a social interaction test that eliminates the possibility of direct physical aggression. We hypothesized that footshock stress would negatively affect sociability, and that those changes might depend on sex and be reversible with the fast-acting benzodiazepine alprazolam. Therefore, we performed social interaction tests 24 hours after two consecutive days of footshock stress. Additionally, we aimed to determine patterns of neuronal activation in the mPFC, CeA, MeA, and ventral ITC associated with social behaviors using expression of the immediate early gene cFos.

## Materials and methods

### Animals

2–4-month-old male and female C57BL/6J mice were obtained from the Jackson Laboratory (Bar Harbor, ME, Stock No: 000664) and housed on a 12 h light/dark cycle with *ad libitum* access to water and chow under standard laboratory conditions. Mice were individually housed for 7 days before the start of and all throughout the experiments. Experiments were performed during the light phase. All animal procedures were performed in accordance with institutional guidelines and were approved by the Institutional Animal Care & Use Committee of Tulane University. Unfamiliar strain-, sex- and age-matched mice were used as the passively interacting counterparts (stimulus mice) during social interaction tests.

### Footshock stress exposure

Footshock exposure or control context exposure was conducted in standard mouse operant conditioning chambers (ENV-307W, Med Associates, Inc., St. Albans, VT) enclosed within sound- and light-attenuating cubicles (ENV-022MD, Med Associates, Inc., St. Albans, VT). The chambers were connected to a computer through an interface and controlled by MED-PC software. The chamber was equipped with a grid floor and a house light, which was cleaned using 70% ethanol.

Seven days after single housing, mice underwent footshock exposure for two consecutive days. Each of the shock sessions included five 1 s, 0.9 mA footshocks presented with a 120 s average pseudorandom intertrial interval (range 90-150 s), totaling 800 s in the chamber. Mice in the control group were exposed to the same chambers for the same period but did not experience footshock.

### Social interaction test

The day after footshock exposure, the mice underwent the social interaction test in a square 46 × 46 × 38 cm arena constructed from sheets of white plexiglass. Behavioral videos were recorded using a digital camera (Allied Vision “Pike” camera, Germany) and Plexon Studio tracking software (Plexon, Dallas, TX). Tests were conducted under dim (10.6 lux) white fluorescent lighting. Stimulus mice were single housed for 3 days before tests. Each of the stimulus mice interacted with three experimental mice with at least 30 min between tests. Experimental mice were perfused 90 min after the test to assess cFos expression.

An indirect social interaction method was chosen to avoid physical aggression between male mice. For the first 3 min, mice were allowed to explore the open arena with two rectangular (15 × 5 × 6 cm) or circular (8 cm diameter, 10 cm high) metallic mesh boxes located in opposing corners 5 cm away from the walls. After the initial exploration, an unfamiliar, untreated stimulus mouse was put underneath one of the boxes. Behavior was recorded for an additional 5 minutes, and sociability was scored using time spent sniffing the mesh box containing the stimulus mouse as a percentage of total box interaction time (mouse preference, %), the latency to approach the stimulus mouse, and the number of defensive tail rattles. Total exploration of the mesh boxes was also scored in seconds to assess general activity. All behavioral measurements were scored by an observer blinded to condition. Consistent with other social interaction scoring protocols [22], sniffing directed to the upper and top part of the mesh boxes, sniffing of feces, bar biting and circulating around the corral without sniffing, were not scored as social interaction.

### Alprazolam treatment

Alprazolam (Sigma-Aldrich, St. Louis, MO) was dissolved in a drop of Tween 80 (Merck, Germany) and saline was added to make a final dose of 0.25 mg/kg. This dose was shown to have anxiolytic effects [23] with minimal motor impairment in C57BL/6J mice [24]. Tween 80 + saline solution was used for vehicle injections. Solutions were administered at 10 ml/kg volume, intraperitoneally, 30 min before social interaction tests.

### Histology

Following testing, mice were anesthetized with 2,2,2-tribromoethanol (240 mg/kg, ip, Sigma) and subsequently transcardially perfused with 4% paraformaldehyde in phosphate-buffered saline (PBS). cFos expression was assessed in mice that were perfused 90 min after the social interaction test. Fixed brains were cut on a compresstome vibrating microtome (Precisionary, Greenville, NC) in 80 μm coronal slices.

Antibody staining was performed on free-floating tissue sections. After 3 × 10 min washes with 0.5% PBST slices were put in 5% donkey serum for 2 hours. Sections were then incubated overnight in primary rabbit anti-cFos antibody (dilution 1:1500; #226 003, Synaptic Systems, Germany) at 4°C. On the next day sections were washed in 0.5% PBST (3 × 10 min), and then went through a 2 hr incubation with secondary donkey anti-rabbit antibody AlexaFluor 488 (dilution 1:500; #A-21206, Thermo Fisher Scientific, Waltham, MA) at 4°C. After 3 × 10 min washes in PBS slices were mounted with mounting medium with DAPI (Biotium, Fremont, CA).

Images were obtained using an AxioScan.Z1 slide-scanning microscope (Zeiss, Germany) and a Nikon A1 Confocal microscope (Nikon, Japan). cFos-positive nuclei were quantified using Fiji ImageJ software (NIH, Bethesda, USA), and averaged for each animal. A blinded observer quantified cFos expression in 2-5 slices per structure per mouse.

### Statistical analysis

Data were analyzed using Prism 9 (GraphPad Software, San Diego, CA). The definition of statistical significance was P ≤ 0.05. Two male mice that showed no interaction with either mesh box during the social interaction test were excluded from analysis. One female outlier that exhibited tail rattling behavior 7 times was also excluded. Because the data were noncontinuous, tail-rattling was analyzed using Fisher’s exact test. To assess the interaction of factors, a 2- or 3-way ANOVA was used. If a significant effect was detected, Sidak’s multiple comparisons test was used, because it assumes independent comparisons and has more power than the Bonferroni method. Correlations between cFos expression and behavior were analyzed for control and stressed mice using pooled data from alprazolam and vehicle treatment groups of both sexes. Using the Kolmogorov-Smirnov test, the data were determined to be non-parametric, so Spearman’s correlation coefficient was used for this analysis. Because there was a Drug x Sex interaction and a main effect of Drug in cFos expression in the IL, we did not include these data in the correlation analysis. For the sake of clarity, we report the results of the interaction tests, the significant simple main effects, and the significant post-hoc tests in the main text. The results of all tests are reported in **Table 1**. All statistical tests were two-tailed.

**Table 1:** Results of Statistical Analyses

| **Effects of alprazolam x footshock stress x sex: 3-way ANOVA** |  |  |  |  |  |
| --- | --- | --- | --- | --- | --- |
| Source | Sum of squares | DFn | DFd | F | p |
| Fig. 1A Sociability (Mouse preference, %) |  |  |  |  |  |
| Alprazolam*Stress*Sex | 0.560 | 1 | 73 | 0.001 | 0.97 |
| Alprazolam*Stress | 2348 | 1 | 73 | 5.71 | **0.02\*** |
| Alprazolam*Sex | 182.4 | 1 | 73 | 0.44 | 0.51 |
| Stress*Sex | 347.5 | 1 | 73 | 0.85 | 0.36 |
| Alprazolam | 490.2 | 1 | 73 | 1.19 | 0.28 |
| Stress | 259.2 | 1 | 73 | 0.63 | 0.43 |
| Sex | 16.89 | 1 | 73 | 0.04 | 0.83 |
| Fig. 1B Latency to approach |  |  |  |  |  |
| Alprazolam*Stress*Sex | 44.45 | 1 | 73 | 0.006 | 0.94 |
| Alprazolam*Stress | 13492 | 1 | 73 | 1.70 | 0.20 |
| Alprazolam*Sex | 12303 | 1 | 73 | 1.55 | 0.22 |
| Stress*Sex | 4270 | 1 | 73 | 0.54 | 0.47 |
| Alprazolam | 478.7 | 1 | 73 | 0.06 | 0.81 |
| Stress | 2646 | 1 | 73 | 0.33 | 0.57 |
| Sex | 11364 | 1 | 73 | 1.43 | 0.24 |
| Fig. 1D Exploration |  |  |  |  |  |
| Alprazolam*Stress*Sex | 47.43 | 1 | 73 | 0.28 | 0.60 |
| Alprazolam*Stress | 257.0 | 1 | 73 | 1.51 | 0.22 |
| Alprazolam*Sex | 190.5 | 1 | 73 | 1.13 | 0.29 |
| Stress*Sex | 0.266 | 1 | 73 | 0.002 | 0.97 |
| Alprazolam | 540.9 | 1 | 73 | 3.20 | 0.08 |
| Stress | 180.2 | 1 | 73 | 1.06 | 0.31 |
| Sex | 3.060 | 1 | 73 | 0.02 | 0.89 |
| **Effects of alprazolam x footshock stress x sex on cFos: 3-way ANOVA** |  |  |  |  |  |
| Fig. 2C IL PFC |  |  |  |  |  |
| Alprazolam*Stress*Sex | 579981 | 1 | 41 | 2.43 | 0.13 |
| Alprazolam*Stress | 172328 | 1 | 41 | 0.72 | 0.40 |
| Alprazolam*Sex | 1378855 | 1 | 41 | 5.78 | **0.02\*** |
| Stress*Sex | 169279 | 1 | 41 | 0.71 | 0.40 |
| Alprazolam | 3014936 | 1 | 41 | 12.6 | **0.001\*** |
| Stress | 2231 | 1 | 41 | 0.01 | 0.92 |
| Sex | 538912 | 1 | 41 | 2.26 | 0.14 |
| Fig. 2D PL PFC |  |  |  |  |  |
| Alprazolam*Stress*Sex | 657.3 | 1 | 33 | 0.008 | 0.93 |
| Alprazolam*Stress | 322417 | 1 | 33 | 3.86 | 0.06 |
| Alprazolam*Sex | 169139 | 1 | 33 | 2.03 | 0.16 |
| Stress*Sex | 7228 | 1 | 33 | 0.09 | 0.77 |
| Alprazolam | 546812 | 1 | 33 | 6.55 | **0.02\*** |
| Stress | 3566 | 1 | 33 | 0.04 | 0.84 |
| Sex | 29411 | 1 | 33 | 0.35 | 0.56 |
| Fig. 2E Capsular CeA |  |  |  |  |  |
| Alprazolam*Stress*Sex | 24040 | 1 | 34 | 0.55 | 0.46 |
| Alprazolam*Stress | 19319 | 1 | 34 | 0.45 | 0.51 |
| Alprazolam*Sex | 54202 | 1 | 34 | 1.25 | 0.27 |
| Stress*Sex | 45335 | 1 | 34 | 1.05 | 0.31 |
| Alprazolam | 38512 | 1 | 34 | 0.89 | 0.35 |
| Stress | 36099 | 1 | 34 | 0.83 | 0.37 |
| Sex | 82985 | 1 | 34 | 1.91 | 0.18 |
| Fig. 2F Medial CeA |  |  |  |  |  |
| Alprazolam*Stress*Sex | 5538 | 1 | 34 | 0.92 | 0.34 |
| Alprazolam*Stress | 3419 | 1 | 34 | 0.57 | 0.46 |
| Alprazolam*Sex | 8969 | 1 | 34 | 1.49 | 0.23 |
| Stress*Sex | 8175 | 1 | 34 | 1.35 | 0.25 |
| Alprazolam | 15566 | 1 | 34 | 2.59 | 0.12 |
| Stress | 307.6 | 1 | 34 | 0.05 | 0.83 |
| Sex | 24836 | 1 | 34 | 4.11 | 0.05 |
| Fig. 2G Lateral CeA |  |  |  |  |  |
| Alprazolam*Stress*Sex | 6744 | 1 | 34 | 0.38 | 0.54 |
| Alprazolam*Stress | 10219 | 1 | 34 | 0.58 | 0.45 |
| Alprazolam*Sex | 762.9 | 1 | 34 | 0.04 | 0.84 |
| Stress*Sex | 11.84 | 1 | 34 | 0.0005 | 0.98 |
| Alprazolam | 11499 | 1 | 34 | 0.65 | 0.42 |
| Stress | 3797 | 1 | 34 | 0.22 | 0.64 |
| Sex | 69608 | 1 | 34 | 3.97 | 0.05 |
| Fig. 2H ITC |  |  |  |  |  |
| Alprazolam*Stress*Sex | 56271 | 1 | 34 | 0.12 | 0.73 |
| Alprazolam*Stress | 46481 | 1 | 34 | 0.10 | 0.75 |
| Alprazolam*Sex | 153940 | 1 | 34 | 0.33 | 0.57 |
| Stress*Sex | 14920 | 1 | 34 | 0.03 | 0.86 |
| Alprazolam | 19359 | 1 | 34 | 0.04 | 0.84 |
| Stress | 9053 | 1 | 34 | 0.02 | 0.89 |
| Sex | 2104455 | 1 | 34 | 4.53 | 0.04* |
| Fig. 2I MeA |  |  |  |  |  |
| Alprazolam*Stress*Sex | 291674 | 1 | 34 | 1.87 | 0.18 |
| Alprazolam*Stress | 4035 | 1 | 34 | 0.03 | 0.87 |
| Alprazolam*Sex | 278969 | 1 | 34 | 1.79 | 0.19 |
| Stress*Sex | 103189 | 1 | 34 | 0.66 | 0.42 |
| Alprazolam | 38692 | 1 | 34 | 0.25 | 0.62 |
| Stress | 6305 | 1 | 34 | 0.04 | 0.84 |
| Sex | 147128 | 1 | 34 | 0.95 | 0.34 |

**Figure 3A: Control group correlations (Spearman's coefficient)**
|  |  | PL | CeC | CeL | CeM | ITC | MeA |
| --- | --- | --- | --- | --- | --- | --- | --- |
| Sociability | r | -.236 | -.627 | -.320 | .118 | -.339 | .313 |
|  | p | .347 | .015* | .244 | .674 | .216 | .255 |
| Exploration | r | .118 | .105 | -.149 | .294 | .375 | .501 |
|  | p | .641 | .706 | .593 | .284 | .168 | .059 |
| Tail rattling | r | .017 | .115 | -.184 | .465 | -.205 | .130 |
|  | p | .947 | .686 | .519 | .086 | .486 | .648 |
| Latency to approach | r | .065 | .557 | .044 | .171 | -.016 | .005 |
|  | p | .797 | .033* | .876 | .538 | .957 | .986 |

**Figure 3B: Footshock stress group correlations (Spearman's coefficient)**
|  |  | PL | CeC | CeL | CeM | ITC | MeA |
| --- | --- | --- | --- | --- | --- | --- | --- |
| Sociability | r | -.663 | -.746 | -.557 | -.307 | -.593 | -.690 |
|  | p | .001** | <.001**** | .007** | .165 | .004** | <.001*** |
| Exploration | r | -.057 | -.310 | -.104 | .034 | -.185 | -.292 |
|  | p | .808 | .161 | .647 | .882 | .410 | .187 |
| Tail rattling | r | -.016 | -.072 | -.209 | -.0149 | -.208 | -.053 |
|  | p | .944 | .750 | .351 | .948 | .353 | .815 |
| Latency to approach | r | .646 | .385 | .245 | -.209 | .399 | .468 |
|  | p | .002** | .077 | .271 | .350 | .066 | .028* |

## Results

### Effects of footshock stress and alprazolam treatment on social behavior

Following two days of footshock stress or context exposure, male (N = 18 control, N = 22 stress) and female (N = 17 control, N = 24 stress) mice were allotted to the vehicle (N = 49, 24 males and 25 females) and alprazolam (N = 32, 16 males and 16 females) treatment groups. These mice were then subjected to a social interaction test designed to measure sociability (**Fig. 1A, B**). A three-way ANOVA revealed that there was statistically significant interaction between the effects of alprazolam and stress on sociability (**Fig. 1C**; drug X stress, F_(1,73)_ = 5.7, *p* = 0.02). Simple main effects analysis showed that there was no statistically significant effect of sex, alprazolam, or stress on sociability (**Fig. 1C; Table 1**). There was no significant interaction between stress, alprazolam, and sex on the latency to approach the social stimulus (**Fig. 1D**; 2-way ANOVA, sex X stress, F _(1, 73)_ = 0.0056, *p* = 0.94). In addition, simple main effects analysis showed that neither stress, alprazolam, nor sex had a significant effect. Further, we applied Fisher’s exact test to analyze noncontinuous tail-rattling data. Interestingly, stressed males displayed significantly more tail-rattling behavior than control males during the social interaction test (*p* = 0.002), while females in both groups displayed equivalent levels of tail rattling (**Fig. 1E**; *p* > 0.99). On the other hand, alprazolam treatment significantly reduced tail-rattling in males (*p* = 0.03) but did not affect tail-rattling in females. In the control vehicle condition, females show higher tail-rattling than males, although this difference did not reach statistical significance (*p* = 0.07). Exploratory behavior, measured as total interaction time with both mesh boxes, was unaffected by footshock stress or alprazolam treatment (**Fig. 1F**; 3-way ANOVA, sex X stress X drug, F _(1, 73)_ = 1.5, *p* = 0.22).

**Figure 1.**
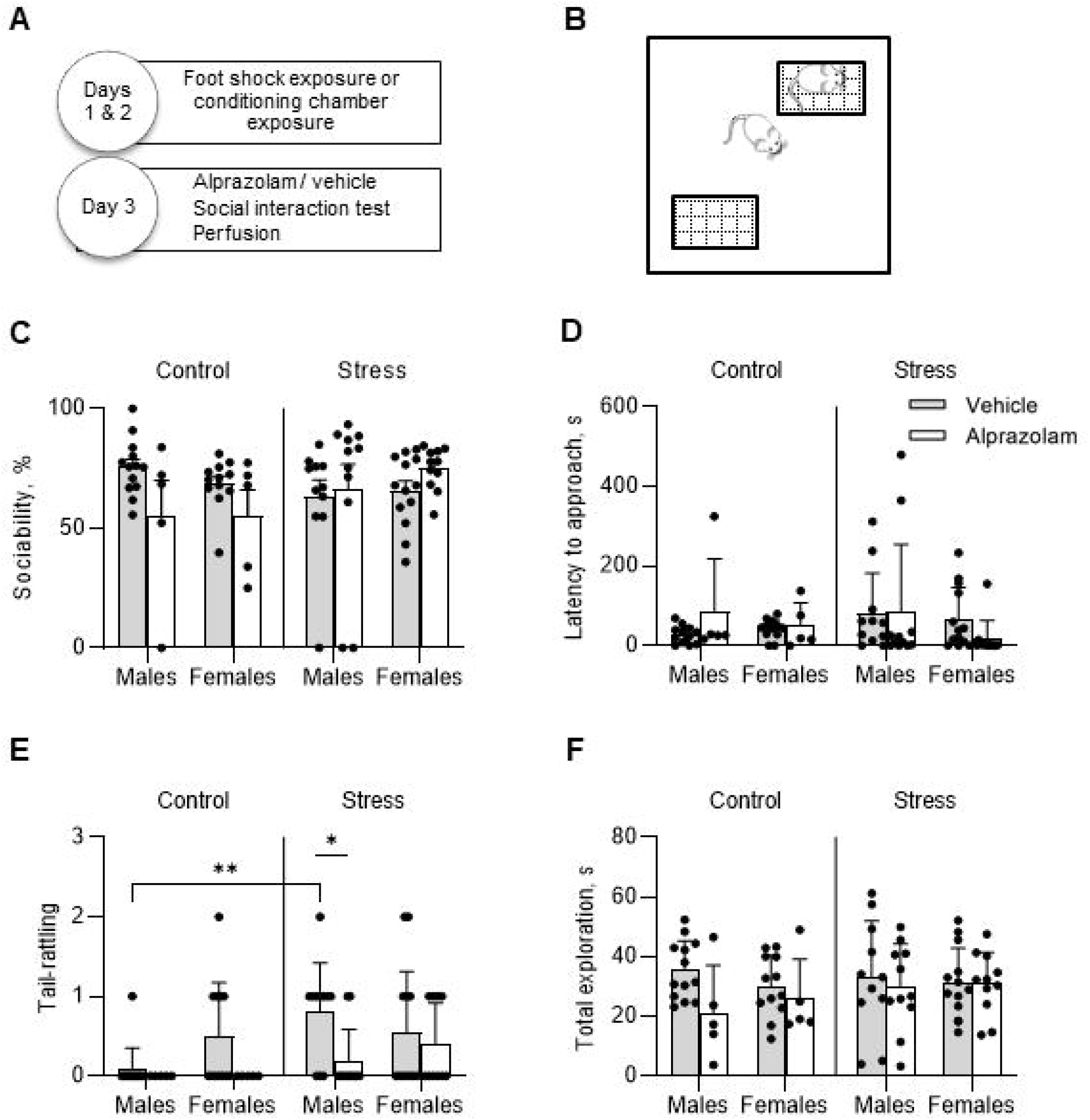
Effects of acute stress and alprazolam on behavior in the social interaction test. **A.** Experimental timeline. **B.** Schematic of the social interaction arena. **C.** There was a significant interaction between the drug and stress condition, but there were no significant results from post hoc multiple comparison tests. **D.** Latency to approach was not significantly affected by stress or alprazolam treatment. **E.** Tail rattling was increased by stress and was significantly reduced by alprazolam in males. **F.** Exploratory behavior was not significantly altered by stress or drug treatment. Data are presented as means□±□SEM. \**p*<0.05 post hoc tests.

### cFos expression analysis

We next quantified expression of cFos in several brain regions involved in the regulation of social behavior (**Fig. 2A, B**). A three-way ANOVA was performed to test for the effects of sex, stress, and alprazolam treatment on cFos expression in the IL, PL, CeA, ventromedial ITC, and MeA (N = 13 control vehicle (5 male, 8 female), N = 10 control alprazolam (5 male, 5 female), N = 14 stress vehicle (5 male, 9 female), N = 12 stress alprazolam (6 male, 6 female)). There was no significant three-way interaction between the effects of these variables on cFos expression in any of the brain areas analyzed (see **Table 1**). There was a significant sex by alprazolam interaction effect on cFos expression in IL (**Fig. 2C**; F_(1,41)_ = 5.78, *p* = 0.02) and a main effects analysis showed that alprazolam treatment significantly reduced cFos expression in the IL (drug effect, F_(1,41)_ = 12.6, *p* = 0.001). Post hoc analysis showed that control females injected with alprazolam had significantly fewer cFos+ cells in the IL compared to vehicle-injected female controls (Sidak’s multiple comparisons test, *p* = 0.03). There was a main effect of drug on cFos expression in the PL (**Fig. 2D**; drug effect, F_(1,33)_ = 6.55, *p* = 0.02). There were no significant simple main effects of sex, stress, or drug on cFos expression in the capsular subdivision of the CeA (**Fig. 2E**), the medial (**Fig. 2F**, sex, F _(1, 34)_ = 4.11, *p* = 0.0505) or lateral subdivision of the CeA (**Fig. 2G**, sex, F _(1, 34)_ = 3.96, *p* = 0.0546), or in the medial amygdala (**Fig. 2I**). There was a significant effect of sex on cFos expression in the ventromedial intercalated nucleus of amygdala, with greater expression levels in males (**Fig. 2H**; sex effect, F_(1,34)_ = 4.53, *p* = 0.04).

**Figure 2.**
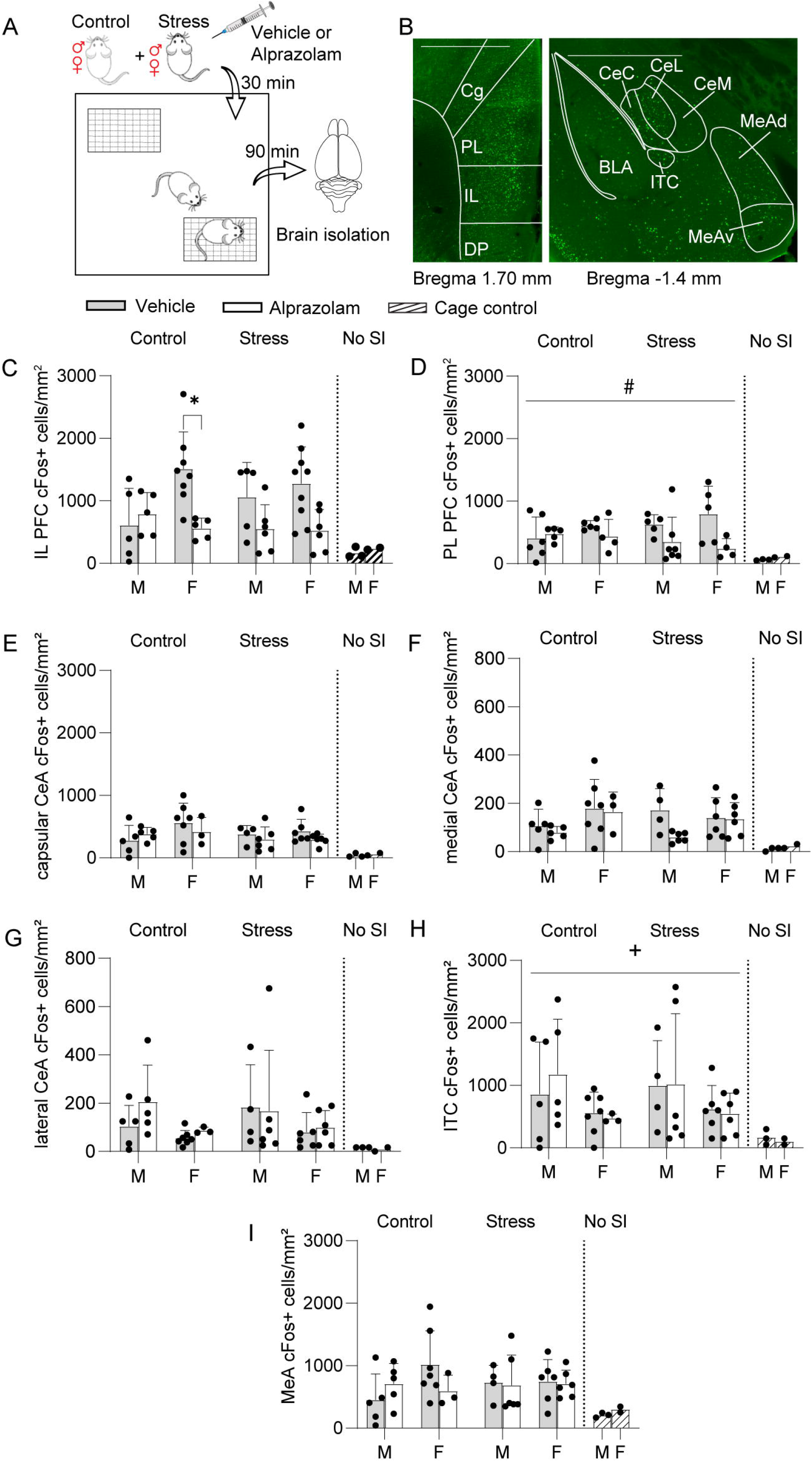
cFos expression patterns following social interaction. **A.** Schematic of the experiment. **B.** Representative images of cFos staining results in mPFC *(left)* and amygdala *(right)*. IL = infralimbic mPFC, CeC = capsular CeA, CeL = lateral CeA, CeM = medial CeA, ITC = ventromedial intercalated nucleus of amygdala, MEAd = dorsal MEA, MEAv = ventral MEA. Scale: 1000 μm. **C-I.** Average number of cFos+ cells / mouse. **C.** Alprazolam reduced cFos expression in the IL in control females. **D**. Alprazolam lowered cFos+ cells in the PL. **E-G.** cFos expression in the capsular, medial, or lateral CeA was not significantly affected by stress, sex, or drug.**H.** cFos expression in the ventromedial ITCs were greater in males. **I.** cFos expression in the MeA was unaffected by stress, sex, or drug. Data are presented as means□±□SEM. \**p*<0.05 post hoc tests. ^+^*p*<0.05, main effect of sex, #*p* < 0.05, main effect of drug.

### Correlations between cFos levels and behavior following the social interaction test

Spearman’s correlation coefficient was used to assess the relationship between behavioral variables and cFos expression levels (**Fig. 3** and **Table 1**). These correlation analyses were performed for the control and stressed groups. For the brain areas in which we did not find any significant effect of drug or sex, we pooled the data from their respective vehicle or alprazolam treatment groups together for analysis. In controls (**Fig. 3A**), there was a significant negative correlation between CeC cFos expression and sociability (r = −0.63, *p* = 0.01), and a positive correlation with latency to approach (r = 0.56, *p* = 0.03). In the stress group (**Fig. 3B**), there was a significant negative correlation between sociability and cFos expression in numerous areas, with more socially active mice demonstrating less neuronal activation in the PL (r = −0.66, *p* = 0.001), capsular (CeC, r = −0.75, *p* = 0.0001) and lateral (r = −0.56, *p* = 0.007) subdivisions of CeA, as well as in the ventromedial ITC (r = −0.59, *p* = 0.003) and medial amygdala (MEA, r = −0.69, *p* = 0.0004).

**Figure 3.**
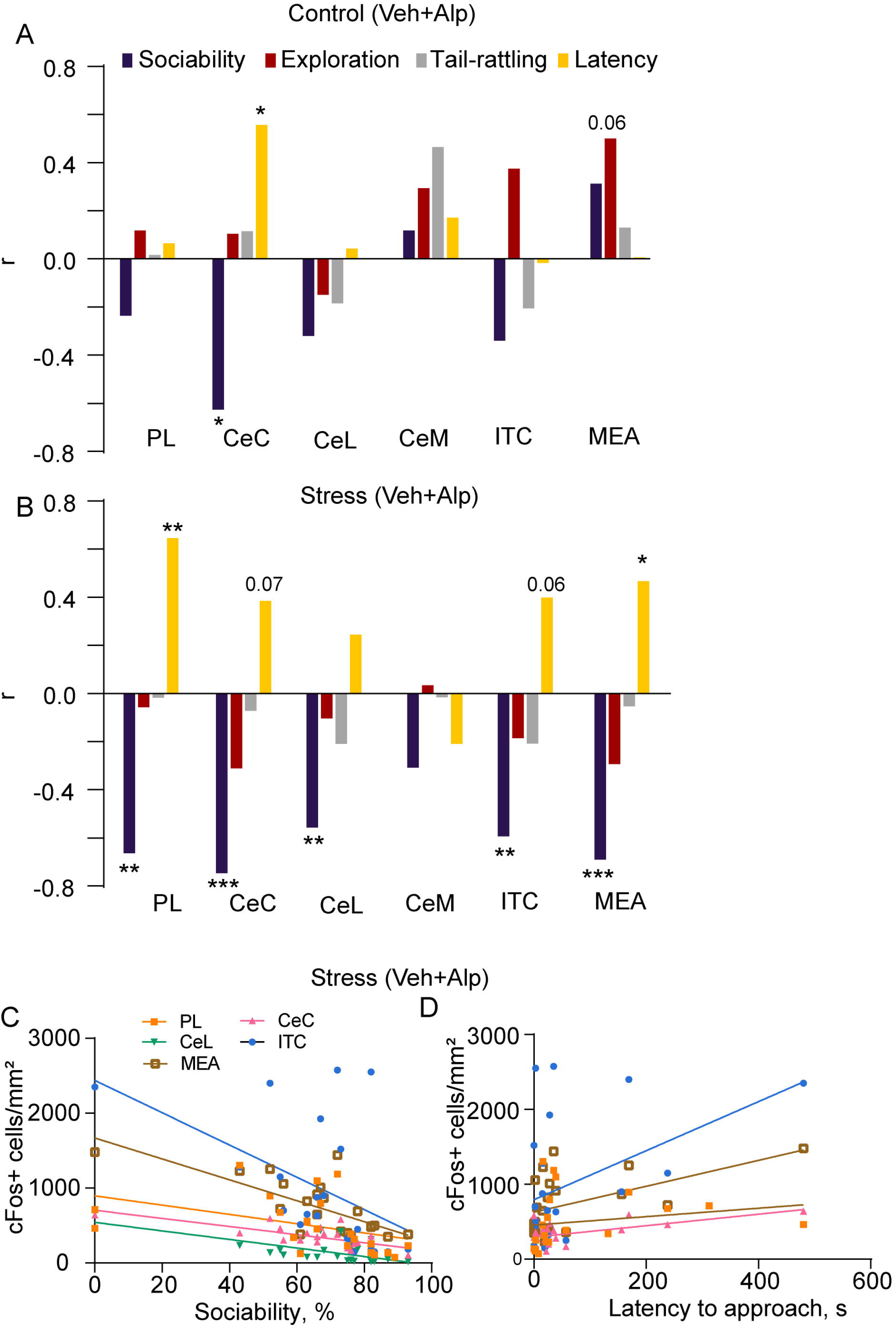
Correlations between behavioral variables and cFos expression. Spearman’s correlation coefficients (r) between the number of cFos+ cells and behavioral parameters in PL, CeA subdivisions, ITC and MeA of vehicle or alprazolam treated mice pooled together in the control (**A**) and stressed (**B**) groups. **C-D**, the distribution of correlation points for sociability and latency to approach against cFos+ cells/mm^2^. Spearman’s correlation coefficient represented with * indicating significant correlations (*p*<0.05). PL = prelimbic cortex, CeC = capsular CeA, CeM = medial CeA, CeL = lateral CeA, ITC = ventromedial intercalated nucleus of amygdala.

Latency to approach the social stimulus positively correlated with the number of cFos+ cells in the PL (r = 0.64, *p* = 0.001), and MEA (r = 0.47, *p* = 0.02), with a trend to significance in

CeC (r = 0.39, *p* = 0.07) and ITC (r = 0.40, *p* = 0.06; **Fig. 3B**). Distribution of data points and regression lines for the respective brain area for sociability and latency is shown in **Fig. 3C-D**.

## Discussion

Exposure to footshock is a commonly used model of acute traumatic stress to assay core features of stress disorders such as social withdrawal [25]; however, sociability after footshock stress has been investigated predominantly in male rodents. In the current study, several measurements of social interaction behavior were taken in male and female mice to investigate potential sex differences in the impact of footshock stress on sociability (**Fig. 1**) and neuronal activation in the mPFC, CeA, and MEA (**Fig. 2**).

Our findings suggest that tail rattling behavior during social interaction is sex-dependent, which is affected by stress specifically in males and diminished by alprazolam (**Fig. 1**). Tail rattling is elicited due to territorial aggression in male mice [26, 27], and it has been suggested that it can be a measurement of threat-induced defensive aggression [27]. Our results are therefore comparable to other mouse sociability studies in which footshock stress produced an enhancement of aggression in male mice [10]. Interestingly, female mice exhibit more tail rattling than males during fear conditioning [28], suggesting that sex differences in tail rattling behavior are both stress- and context-dependent. The significant reduction of tail rattling by alprazolam further supports the link between this behavior and negative valence states. Overall, the observed sex differences reinforce the idea that sex is a crucial factor that should be considered in stress-related studies.

We found a significant interaction between the effects of alprazolam and stress on preference for the social stimulus with alprazolam treatment leading to a decrease in sociability in the control group, while there were no such trends in the stress group. There is evidence that social interaction itself is anxiogenic, even under control conditions [29]. This is further supported by the negative correlation between sociability and cFos expression in the CeA (**Fig. 3**). Acute stress by itself did not significantly lower sociability in contrast to other studies that were focused on the effects of stress on social behavior, which mostly employed chronic stress models [7]. Because the mice in our experiment underwent 9 days of single housing before the social interaction test, it could be hypothesized that social isolation could enhance social interest, thus leveling out social withdrawal [30].

Our results show that cFos expression in the ITC is higher in males compared to females after social interaction. Given the known role of amygdala regions in mediating threat responses [17, 31], these data suggest that social interaction may induce a higher level of defensiveness in male mice. Consistent with this hypothesis, previous work has shown that CeA neuronal activation is associated with male mouse aggression during social interaction in the resident intruder assay [12, 32]. These findings emphasize the need to study sex differences while deciphering the relationship between stress and social behaviors. Further investigations should be conducted into sex differences in different neuronal populations of CeA, expressing molecular markers such as somatostatin or corticotropin-releasing factor. This is especially relevant given that there is a greater number of corticotropin-releasing factor receptor 1-containing neurons in the male compared to female CeA, and they have different neuronal excitability in response to corticotropin-releasing factor [33].

IL and PL cFos levels were decreased due to alprazolam treatment both in stressed and control mice. Similarly, in humans, benzodiazepines decrease cerebral blood flow in PFC [34]. cFos expression in PL and IL mPFC in our study was not significantly affected by stress. Thus, although the IL has been shown to be activated in response to a social cue [9], the mPFC does not seem to modulate the interaction between traumatic stress and sociability in mice. Acute restraint stress has been shown to hyperactivate PL, but not IL or CeA, and social interaction following stress further activated PL PFC and did not affect total CeA activation [35]. A similar lack of differences in cFos expression was observed in our study, which may occur due to the delay between stress exposure and sacrifice of the animals.

Among control vehicle-injected mice, females had more cFos+ cells in IL. Similar results were obtained by Tan et al. who observed that pyramidal neurons in the mPFC of females after chronic adolescent social isolation stress showed a blunted increase of discharge rates during sociability tests [36]. Sex differences in the modulation of social behavior by the mPFC were previously explored regarding oxytocin-receptor-expressing interneurons which have differential responses to oxytocin in male and female mice and regulate the social motivation of female mice to interact with male mice during estrus [37]. Alprazolam significantly lowered the number of cFos+ cells in the IL compared to vehicle-injected female controls, consistent with the benzodiazepine effects observed in humans: women demonstrate decreased activity in frontal regions after treatment, while an opposite effect is present in males [38]. This effect could be related to females having significantly higher GABA-A benzodiazepine receptor availability [39].

The MEA has been linked to a wide variety of social behaviors, such as aggression, mating, and parenting [40]. Here, we find that MEA cFos expression induced by the social interaction test is not affected by sex, stress, or alprazolam. Prior studies have demonstrated that socially defeated females housed with aggressive male residents exhibit increased cFos activation in the medial amygdala [41], and the MEA is more responsive to aggressive than to benign social interaction [42]. We designed our experiments to exclude the possibility of aggressive interactions, therefore free interactions between the experimental and stimulus mice were not possible. Future studies could investigate the effects of stress and sex on MEA activation in free interaction paradigms.

In stressed animals, sociability was significantly negatively correlated with cFos expression in the PL, CeC and CeL, as well as in the ventromedial ITC and MEA. In non-stressed animals, MEA activation has been demonstrated previously as a result of social interaction [43], and an opposite relationship has been shown between CeA cFos expression and sociability after anxiogenic synthetic amphetamine treatment [44]. Because mPFC activation has been demonstrated to suppress social behaviors [45], the negative correlation observed in the current study could be expected, although enhanced activity of a subset of mPFC neurons was correlated with social approach behavior previously [46]. In controls, there was a significant negative correlation between sociability and CeC cFos expression, and an opposite correlation between the latter and the latency to approach. Latency to approach the social stimulus also positively correlated with the number of cFos+ cells in the PL and MEA.

The social consequences of stress have been extensively studied using the social defeat stress model that entails exposure to emotional / psychological stress and leads to depression-related outcomes [7]. After the social defeat procedure, most mice develop a decreased drive to approach and interact with the social target [47]; however, social defeat stress is difficult to achieve in female mice. Acute stress paradigms utilizing footshock facilitate investigations into sex differences in social behavior following trauma. Future work should define the optimal conditions, such as footshock intensity or lighting conditions, that influence sociability after stress. It could be especially valuable to develop paradigms that stratify mice as resilient and susceptible to further validate acute footshock stress as a tool for PTSD research.

## Author contributions

Conceptualization—MD, CDB, JPF

Formal analysis—MD, CDB, JPF

Funding acquisition—JPF

Investigation—MD, CDB, LSO, KW, AR, SB, EB, AD, AS

Methodology—MD, CDB, JPF

Project Administration / Supervision—MD, JPF

Resources—JPF

Visualization—MD, CDB, JPF

Writing, original draft—MD, JPF

Writing, review & editing—MD, JPF, CDB, LSO, KW

